# Cannabidiol Attenuates Pulmonary Arterial Hypertension by Normalizing the Mitochondrial Function in Vascular Smooth Muscle Cells

**DOI:** 10.1101/2020.11.04.364547

**Authors:** Xiaohui Lu, Jingyuan Zhang, Huijiao Liu, Wenqiang Ma, Leo Yu, Xin Tan, Shubin Wang, Fazheng Ren, Xiru Li, Xiangdong Li

## Abstract

Pulmonary artery hypertension (PAH) is a chronic disease associated with enhanced proliferation of pulmonary artery smooth muscle cells (PASMCs) and dysfunctional mitochondria, which was with limited therapeutic options. It has been proved that cannabidiol (CBD) had antioxidant effects in many cardiovascular diseases, whereas the efficacy of CBD in PAH is unknown. To defined the effect of CBD in PAH, we explored the functions of CBD in both PASMCs proliferation test in vitro, and preventive and therapeutic PAH rodent models in vivo. The roles of CBD in mitochondria function and the oxidant stress were assessed in human PASMCs and PAH mice. We found that CBD significantly inhibited hyperproliferation of hypoxia-induced PASMCs, and intragastrically administered CBD could reverse the pathological changes in both Sugen-hypoxia and MCT-induced PAH mice models. Mechanical analysis demonstrated that CBD alleviated PAH by recovering mitochondrial energy metabolism, normalizing the hypoxia-induced oxidant stress, inhibiting abnormal glycolysis and lactate accumulation in cannabinoids receptors-independent manner. Thus, CBD could be a potential drug for PAH.

**Graphical Abstracts:** 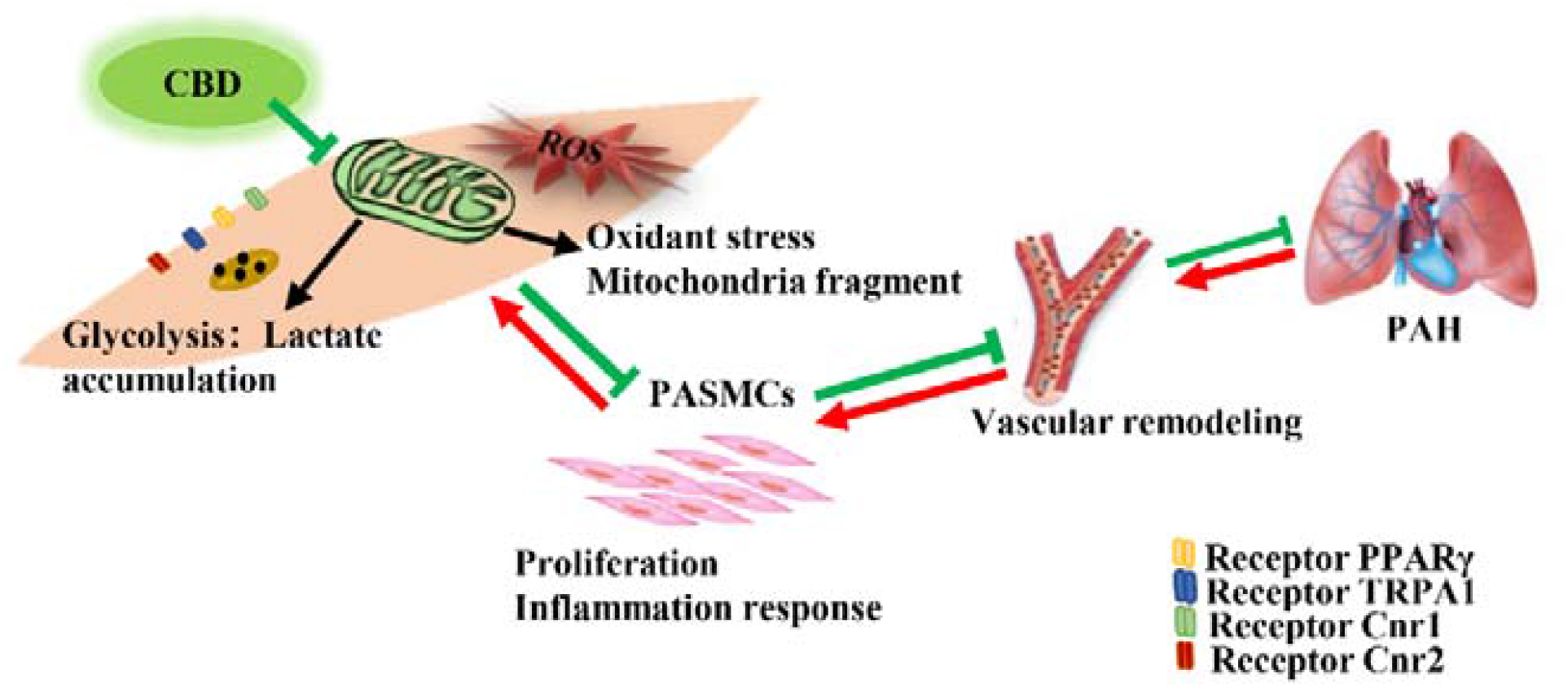

## Introduction

Pulmonary artery hypertension (PAH) is a fatal disease associated with vascular cells dysfunction, proliferation and inflammatory infiltration[1, 2]. Growing evidences have implicated that mitochondria were involved in the pathogenic mechanism in PAH [3–5]. Mitochondrial fission and fusion play critical roles in maintaining functional mitochondria when cells undergo metabolic or environmental stresses [6]. Furthermore, PAH is closely associated with the change of oxidant state intracellular, and there are many antioxidant factors maintaining vascular homeostasis, such as the nuclear factor E2 related factor 2 (NFE2L2), heme oxygenase 1 (HMOX-1), quinone oxidoreductase 1 (NQO1) and superoxide dismutase (SOD), whose expression levels or enzyme activities can partially reflect the oxidation state of the organism [7, 8]. Cells under inflammation preferentially use glycolysis as a source of energy, whereas during the resolution phase they rely mainly on OXPHOS metabolism [9]. A shift from oxidative phosphorylation to glycolysis occurred in PAH pulmonary vessels and the right ventricle [10], these two respiration methods maintain the metabolic balance in the cells of the organism.

At present, five classes of drugs targeting PAH, such as bosentan, sildenafil, riociguat, epoprostenol and selexipag, have been approved by FDA [11]. Even though great advances have been made in PAH treatment, patients still suffer from the disease progression and severe side effects like liver toxicity. New drugs with high efficacy and less side-effect are urgently needed to be explored.

*Cannabis sativa* is a remarkable plant containing many valuable natural components, consisted of the 483 known Phyto-compounds used for medical or recreational purposes, including more than 60 unique compounds known as cannabinoids[12, 13]. Except the primary psychoactive constituent delta-9-tetrahydrocannabinol (THC), the non-psychoactive constituents cannabidiol (CBD), tetrahydrocannabivarin (THCV), cannabidivarin (CBDV) and cannflavin A (CFA) have been widely reported to elicit many therapeutic effects in analgesia, anti-inflammatory and so on [14]. More recently, CBD had been approved by FDA in 2018 for reducing seizures related to a rare form of pediatric epilepsy [15]. Studies suggested that CBD had therapeutic utility in pathological conditions of heart dysfunction and vascular abnormality, it improved both heart and arterial performance with its anti-inflammatory and antioxidant effects[16, 17], whereas the effect of CBD in PAH is still unknown.

Here, we wanted to clarify the therapeutic effect of CBD in the development of PAH in the current study.

## Materials and Methods

The data that support this study are available from the corresponding author upon reasonable request. Extended Material and Methods section are available in the Supplementary material online [18–30].

### Statistical Analysis

All graphs were performed using GraphPad Prism 8.0.2 (GraphPad Software, USA). The data was analyzed by SPSS12.0.1 (SPSS, USA). All results were shown as mean ± SEM. Comparisons of means between 2 groups were performed by unpaired Student’s t-test for normally distributed cases. Comparisons of means among three or more groups were performed by one-way or two-way ANOVA (analysis of variance) followed by Bonferroni’s multiple comparison test or the Dunnett’s method for multiple comparison, as appropriate. All reported *P* values were 2-tailed, with a *P* value of less than 0.05 indicating statistical significance.

## Results

### 3.1 CBD inhibited mice PASMCs proliferation without cytotoxicity

To find a novel drug for PAH patients, several cannabinoid compounds extracted from *Cannabis sativa*, including CBD, CBDV, THCV, CFA, have been screened in mice PASMCs. Previously, our results showed that CBDV, THCV, CFA did not inhibit the hyperproliferation of mice PASMCs, and some of them were with various degrees of cytotoxicity. Thus, we focused on CBD, which exhibited strong anti-proliferation of mice PASMCs and less toxicity. The purity of the isolated mice PASMCs was confirmed above 90% by IHC (Figure 1A). We found that CBD at 10 μM reduced the expression of *Il6* with no effects on cell viability (Figure 1B and 1C). Normally, the lactate dehydrogenase (LDH) is present in the living cells and will be leak out once the cells die, and it can be used for estimating the cytotoxicity. In our study, CBD at 20 μM showed higher cytotoxicity and reduced cell viability in mice PASMCs (Figure 1D, 1E). So, the better concentration of CBD on mice PASMCs was 10 μM. And the cell proliferation assay confirmed that CBD at 10 μM could inhibit the hyperproliferation of hypoxia-induced mice PASMCs (Figure 1F-G).

**Figure 1.**
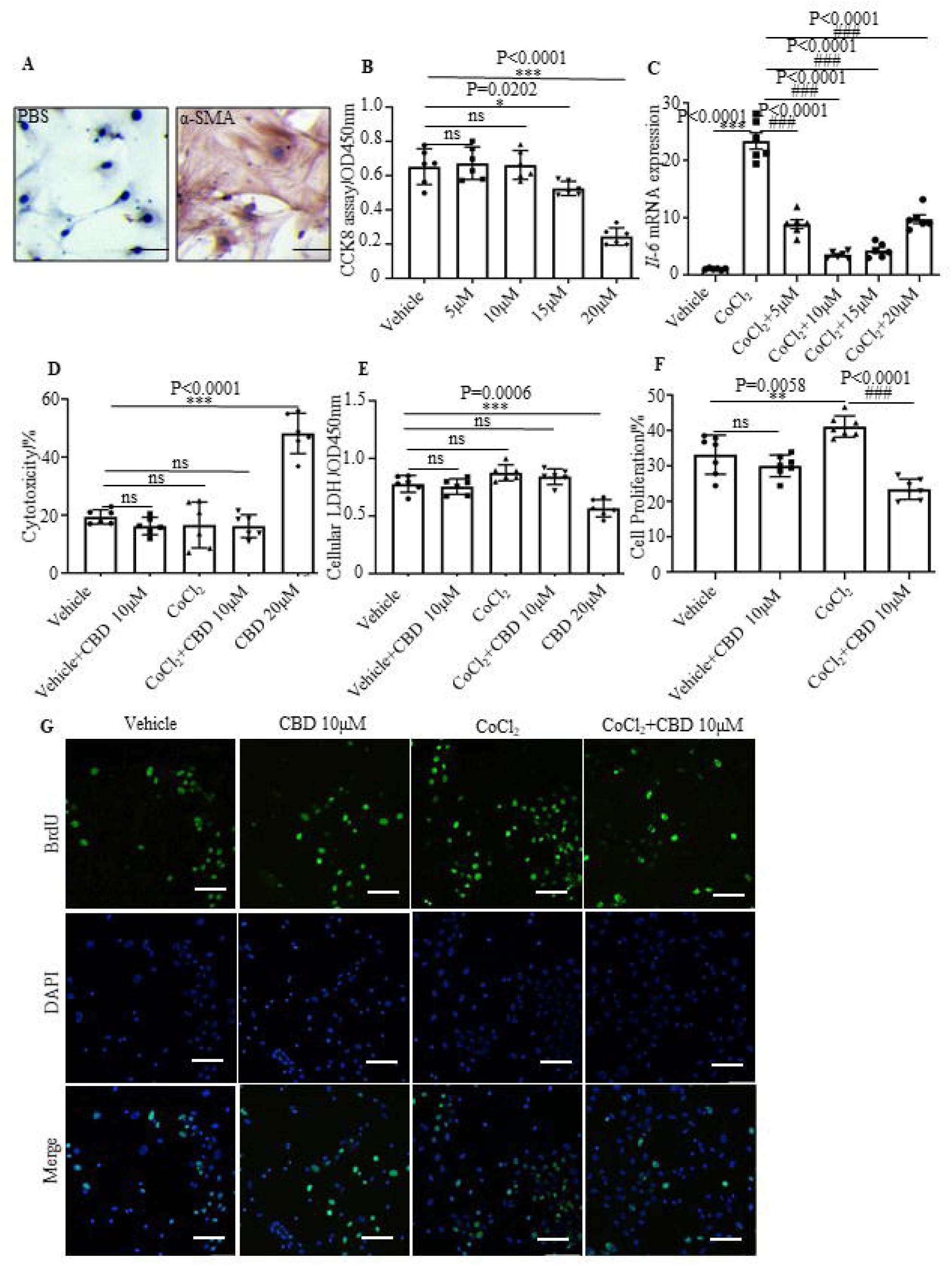
Inhibition of CBD on mice PASMCs hyperproliferation without cytotoxicity. **A**, the purity of mice PASMCs assessed by IHC with the α-SMA antibody. **B**, Cell viability assessed by CCK8 assay, with concentration of CBD at 5 µM, 10 µM, 15 µM, 20 µM (n=6). **C**, Expression level of *Il6* in the mice PASMCs, incubation with different concentration of CBD and/or CoCl_2_ at 200 µM for stimulating the hypoxia condition (n=6). **D** and **E**, the level of LDH was assessed with LDH detection assay in both extracellular (cytotoxicity) and intracellular (cell viability) (n=6). scale bar=100 µm. **F**, Quantitative assessment of BrdU antibody to calculated the ratio of PASMCs proliferation. **G**, Immunofluorescence of BrdU positive ratio of the PASMCs by Fluorescence microplate reader (n=7), the nuclei of cell were stained with DAPI. Mean +/− SEM. The results were analyzed by one-way ANOVA followed by Bonferroni’s multiple comparison test, \**P* < 0.05, \*\**P*< 0.01, \*\*\**P* < 0.001 vs. the control group, and #*P*< 0.05, ##*P*< 0.01, ###*P* < 0.001 vs. the CoCl_2_ group.

### 3.2 CBD reversed the phenotypes in two conventional PAH mice models

To explore the efficacy of CBD in the PAH, we performed two sets of experiments in vivo, including Sugen (SU-5416) hypoxia- and monocrotaline (MCT)-induced PAH models (Set □ for prevention model, Set □ for therapeutic model, respectively). In Set □, we used the Sugen hypoxia-induced and MCT-induced PAH preventive mice models, mice treated with CBD at 10 mg/kg showed attenuated PAH phenotypes, compared with the hypoxia-induced mice and the MCT-induced mice, including the reductions in right ventricular systolic pressure (RVSP), right ventricular hypertrophy (RVH) (Figure 2A-B, Figure S1A-B), the PAH-related inflammation and the remodeling rate of the pulmonary arteries (Figure 2C-F, Figure S1C-F).

**Figure 2.**
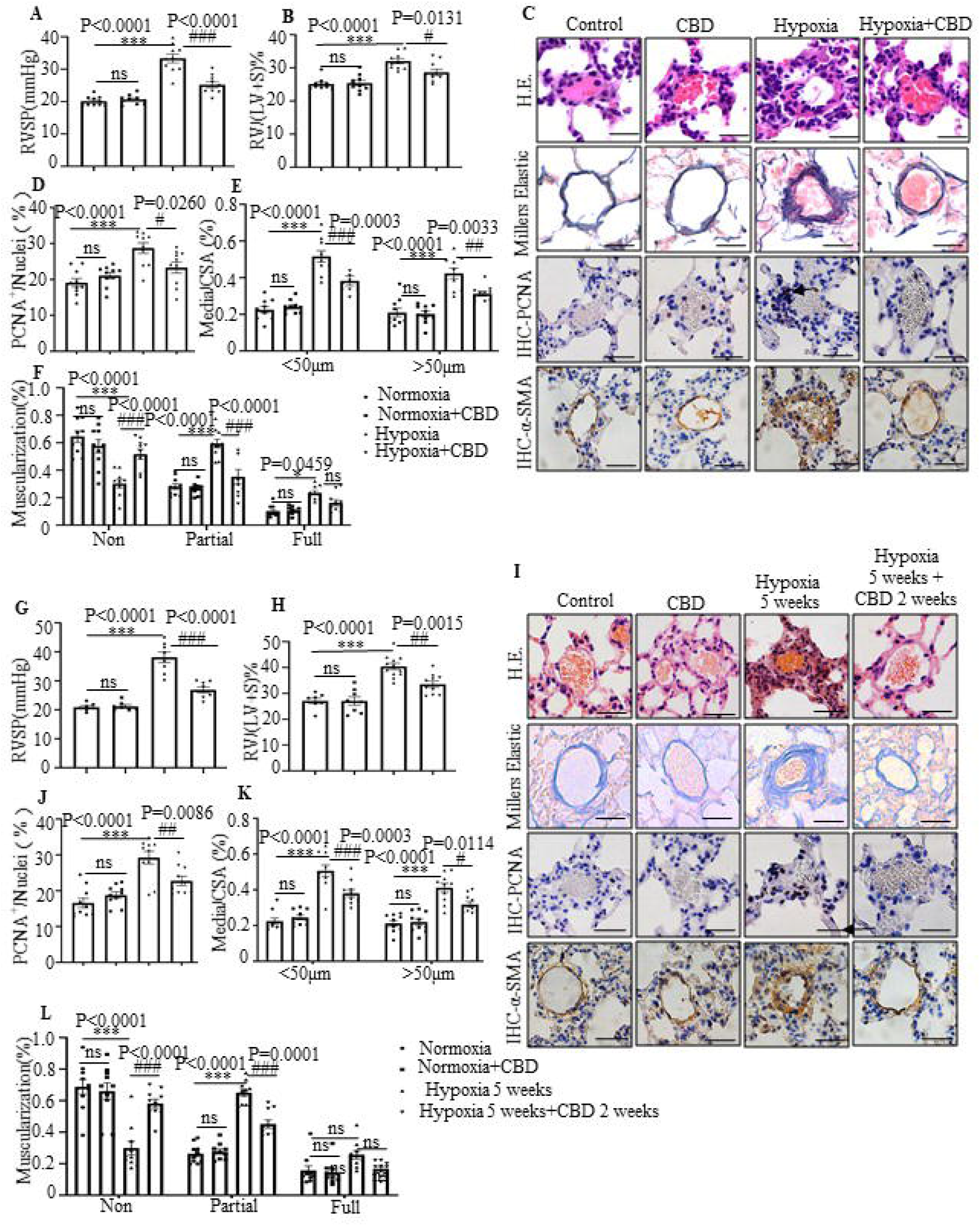
CBD reversed pathological progression of Sugen-hypoxia induced PAH preventive and therapeutic mice models. **A-F**, The data from PAH preventive mice model. **G-L**, The data from PAH therapeutic mice model. **A** and **G**, Assessments of RVSP. **B** and **H**, Assessments of RVH. **C** and **I**, Representative images (×400 magnification) of pulmonary arteries stained with H&E, and representative images (×400 magnification) of vascular remodeling in the distal arterioles stained with elastin or immunostained for PCNA and α-SMA. **D-F** and **J-L**, Pulmonary vascular remodeling rate in the Sugen-hypoxia mice, including the quantification of the relative number of PCNA^+^/nuclei, the degree of medial wall thickness as a ratio of total vessel size (Media/CSA), and the proportion of non-, partially-, or fully-muscularized pulmonary arterioles (25 to 75 μm in diameter) from PAH model mice (n = 10 mice per group). scale bar=20 µm. Mean +/− SEM. n=10. The results were analyzed by one-way ANOVA followed by Bonferroni’s multiple comparison test, \**P*< 0.05, \*\**P*< 0.01, \*\*\**P* < 0.001 vs. the control group, and #*P*< 0.05, ##*P*< 0.01, ###*P* < 0.001 vs. the hypoxia group.

In Set II, we examined the efficacy of CBD in the established Sugen hypoxia-induced and MCT-induced PAH therapeutic mice models, these mice were characterized with higher RVSP and RVH (Figure 2G-H and S2A-B) as well as extensive pulmonary arterial muscularization (Figure 2I-L and S2C-F). After daily administrated with CBD (10 mg/kg, i.g.) for 2 weeks, we obtained that attenuated lung phenotypes in PAH mice, including the reductions in RVSP, RVH and vascular remodeling, compared to saline treated controls (Figure 2 and S2).

Regarding to the strong inhibition of CBD in PAH, we compared it with the compounds used in the first-line therapy against PAH, such as Bosentan and Beraprost Sodium. The results illustrated that all these three compounds attenuated the elevated RVSP and RVH in the Sugen hypoxia-induced PAH mice, and the efficacies among CBD, Bosentan and Beraprost Sodium were no significant differences in the prevention of Sugen-hypoxia induced PAH (Figure S3A-B).

### 3.3 CBD inhibited inflammation independent of the cannabinoid’s receptors in PAH

CBD treatment significantly suppressed the expression levels of *Il6* and *Tnf*α in the mice lung from the Sugen-hypoxia and MCT-induced models, as well as the mRNA expression levels of chemokines *Ccl2* and *Cxcl10* in mice PASMCs (Figure 3A-C). As CBD has been reported to elicit its molecular action via cannabinoids receptor 1 (Cnr1), cannabinoids receptor 1 (Cnr2), transient receptor potential A1 (TRPA1), peroxisome proliferator-activated receptor (PPARγ) receptors [31], we then assessed the expression level of *Il6* in mice PASMCs with antagonists or channel blockers for receptors of cannabinoids, such as rimonabant, SR144528, AM630, GW9662 and HC030031 (Figure 3D-F and Figure S4). The results showed that CBD could normalize hypoxia-induced expression of *Il6*, but antagonists of Cnr1, TRPA1 and PPARγ did not counteract the anti-inflammatory effect of CBD in mice PASMCs, while the expression of *Il6* in the PASMCs treated with Cnr2 antagonists (SR144528, AM630) (in combination with CBD under hypoxia condition) was increased compared to the CBD treatment alone under hypoxia condition (Figure 3D-E). To clarify whether the inhibition of CBD on PAH was mainly via Cnr2 signal pathway, we generated the Cnr2^-/-^ mice. In Sugen-hypoxia induced Cnr2^-/-^ PAH mice models, we observed no significant phenotypical and pathological differences between the wild type PAH mice and the Cnr2^-/-^ PAH mice. However, CBD at 10 mg/kg can predominantly reverse the phenotypes of PAH in both wild type mice and Cnr2^-/-^ mice, including RVSP, RVH, and pathological index of PAH (Figure 3G-I). All these results indicated that Cnr2 was not involved in the inhibition of CBD in PAH development.

**Figure 3.**
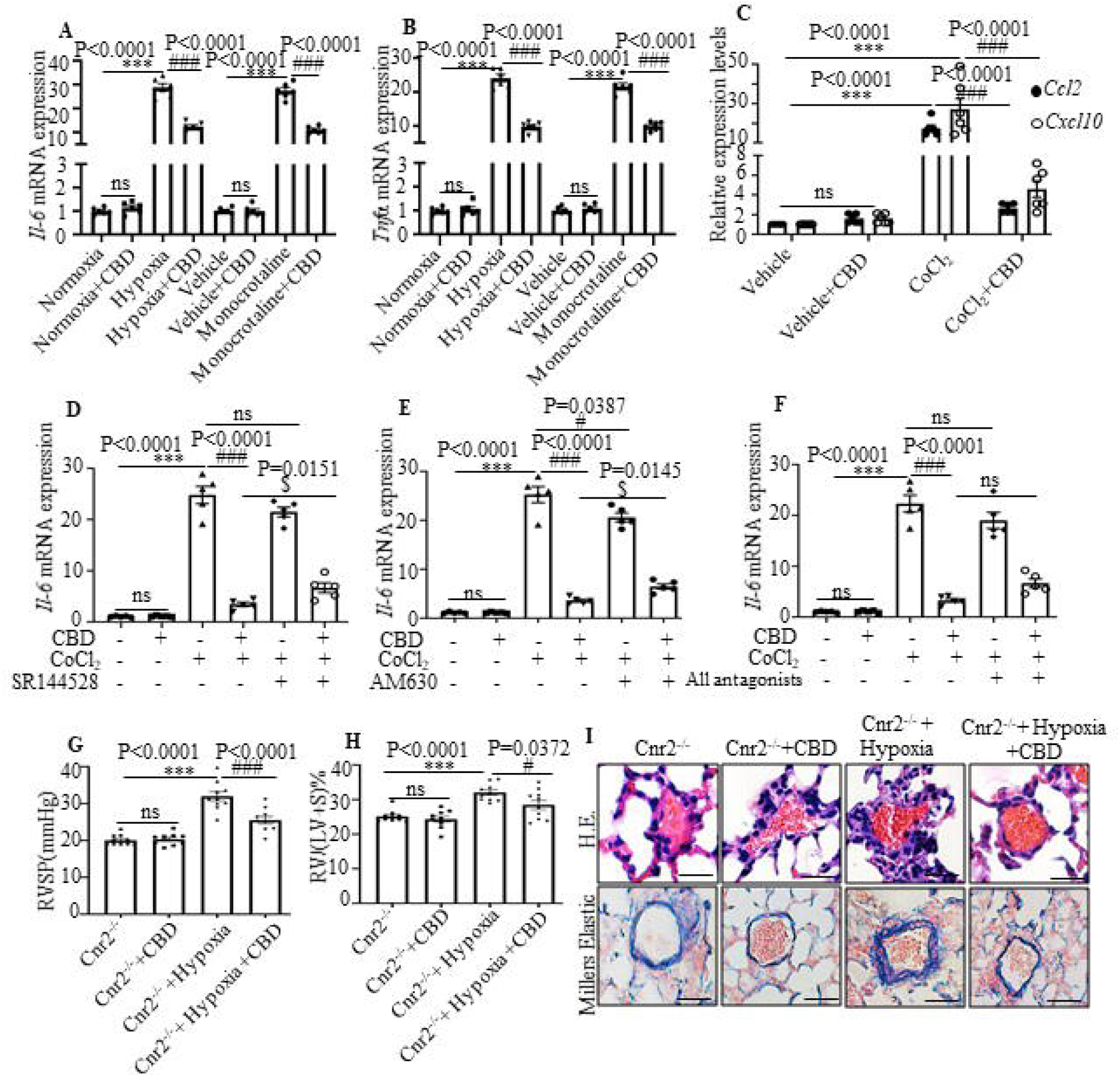
CBD inhibited inflammation without canonical cannabinoids receptors involvement in PAH. **A-B**, Expression of *Il6* and *Tnf*α mRNA assessed in the lung of hypoxia- and MCT-induced PAH mice models. **C**, Expression of *Ccl2* and *Cxcl10* were assessed in the mice PASMCs with or without CoCl_2_ treatment. **D-F**, Expression of *Il6* treated by CoCl_2_, and the effect of antagonist or channel blocker of CBD receptors (SR144528, AM630 and both) in mice PASMCs, the concentration of them were 10 µM, which were equal to the concentration of CBD, CBD were pre-treated for 30 min with either the antagonists or inhibitors. n=6. **G-H**, Assessments of RVSP, RVH and pulmonary vascular remodeling rate in the PAH Cnr2^-/-^ mice models, n=10 each group. **I**, Representative images (×400 magnification) of pulmonary arteries stained with H&E and elastin at 6-month-old mice, Cnr2^-/-^ and WT littermate controls with or without hypoxia-induced with subcutaneous injections of Sugen for 21 days and daily CBD-treated i.g., scale bar=50 µm. The results were analyzed by one-way ANOVA followed by Bonferroni’s multiple comparison test, \**P* < 0.05, \*\**P*< 0.01, \*\*\**P* < 0.001 vs. the control group, and #*P* < 0.05, ##*P*< 0.01, ###*P* < 0.001 vs. the hypoxia or MCT, or CoCl_2_ treatment group, $*P*<0.05 vs. the CoCl_2_ with CBD group.

### 3.4 CBD restored mitochondrial function and oxidant stress in human PASMCs and PAH mice

By using MitoTracker, we found that CBD treatment could normalize the malfunctioned mitochondrial networks (increased the proportion of lineage that longer than 1 µm and mean length of mitochondria/pixels of the mitochondrial networks) in hypoxia-induced human PASMCs (Figure 4A-C). Next, we detected a set of genes that reported to promote mitochondrial fission, including dynamin-1-like protein (*DRP1*), mitochondrial fission 1 protein (*FIS1*) and optic atrophy type 1 (*OPA1*), and a group of genes for mitochondrial fusion, such as mitofusin 1 (*MFN1*), mitofusin 2 (*MFN2*), and mitochondrial elongation factor (*MIEF1*) [32]. We found that CBD treatment significantly reduced the expression levels of *DRP1, FIS1, OPA1*, whereas hypoxia upregulated the expressions of these genes in human PASMCs (Figure 4D-F). Conversely, CBD increased the expressions of *MFN1, MFN2*, and *MIEF1* in human PASMCs (Figure 4G-I). The protein level of MFN2 was consist with the RNA expression (Figure S5J). Together, CBD normalized the mitochondria morphology of the hypoxia-induced human PASMCs and sustained the cell homeostasis.

**Figure 4.**
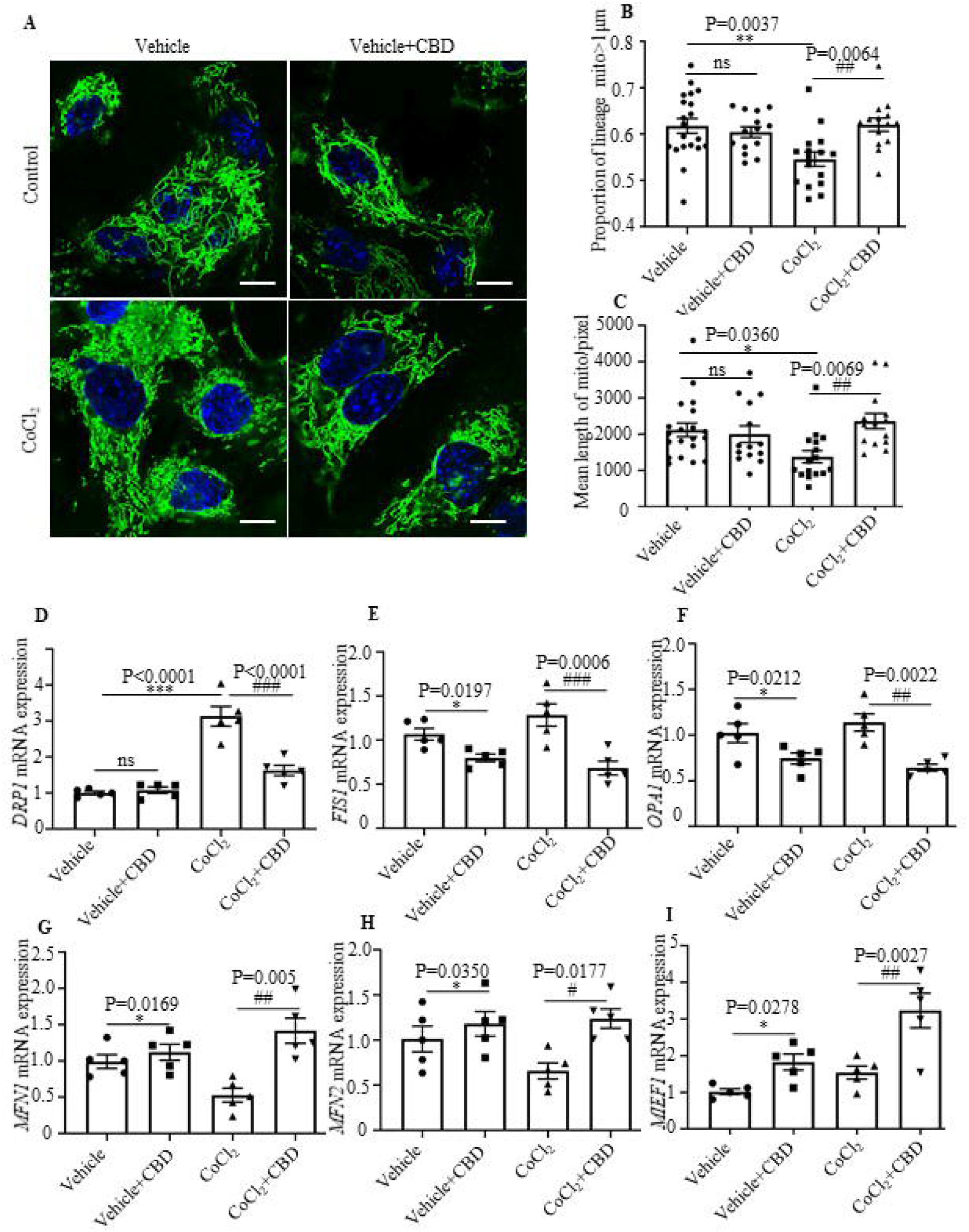
CBD normalized mitochondrial function in hypoxia-induced PASMCs. **A**, Representative images of control human PASMCs and CoCl_2_-treated human PASMCs labeled with MitoTracker after the treatment with CBD (10 µM) or the control vehicle for 2 h. Nuclei were counterstained using Hoechst 33342, Scale bars=10µm. **B** and **C**, Quantification of mitochondrial proportion of lineage that longer than 1 µm (40 pixels, about 1µm) and mean length of mitochondria/pixels in control human PASMCs and CoCl_2_-treated human PASMCs labeled for mitochondria after the treatment with CBD or the control vehicle for 2 h. **D**-**I**, RT-PCR analysis of *DRP1, FIS1, OPA1, MFN1, MFN2, MIEF1* in CoCl_2_ treated human PASMCs after the treatment with CBD (10 µM) or vehicle for 12 h. n=6. Mean +/− SEM. The results were analyzed by one-way ANOVA followed by Bonferroni’s multiple comparison test, \**P* < 0.05, \*\**P*< 0.01, \*\*\**P* < 0.001 vs. the control group, and #*P* < 0.05, ##*P*< 0.01, ###*P* < 0.001 vs. the CoCl_2_ treatment group.

CBD has been reported to affect redox balance by modifying the levels and activities of both oxidants and antioxidants [33, 34]. As we all known, glutathione reductase (GR) and glutathione peroxidase (GSH) protected the structure and function of cell membranes from peroxides damage [35]. We observed that CBD could normalize the activities of GR and GSH in the blood of PAH mice (Figure 5A-B). The content of malondialdehyde (MDA), a product of peroxidation, was elevated in the blood of PAH mice, and this elevation could be significantly reduced by CBD treatment (Figure 5C). Furthermore, to assess the antioxidant function of CBD, the mRNA expressions of some canonical genes in the signal pathway of oxidative stress were examined. We observed that CBD significantly reduced the expression level of *Keap1* (Kelch-like ECH-associated protein 1) in mice PASMCs. Consistently, CBD upregulated the expressions of *Nrf2* (*Nfe2l2*) (a master regulator of cellular response against oxidative stress), and its downstream genes, such as, *Nqo1, Hmox1*, and *Sod1* compared to the hypoxia-induced mice PASMCs (Figure 5D-H). Moreover, we observed that CBD significantly normalized the high levels of cytosolic ROS in hypoxia-induced human PASMCs compared with the control PASMCs (Figure 5I-J). Furthermore, by using Seahorse XF24-3 apparatus, we evaluated the effects of CBD treatment on the O_2_ consumption rate (OCR) and extracellular acidification rate (ECAR) of human PASMCs. We observed significantly decreased production of ATP, maximal respiration, and OCR/ECAR ratio in hypoxia-induced human PASMCs compared with the control PASMCs, whereas these parameters were significantly normalized by CBD treatment (Figure 6A-B and Figure S5A-D).

**Figure 5.**
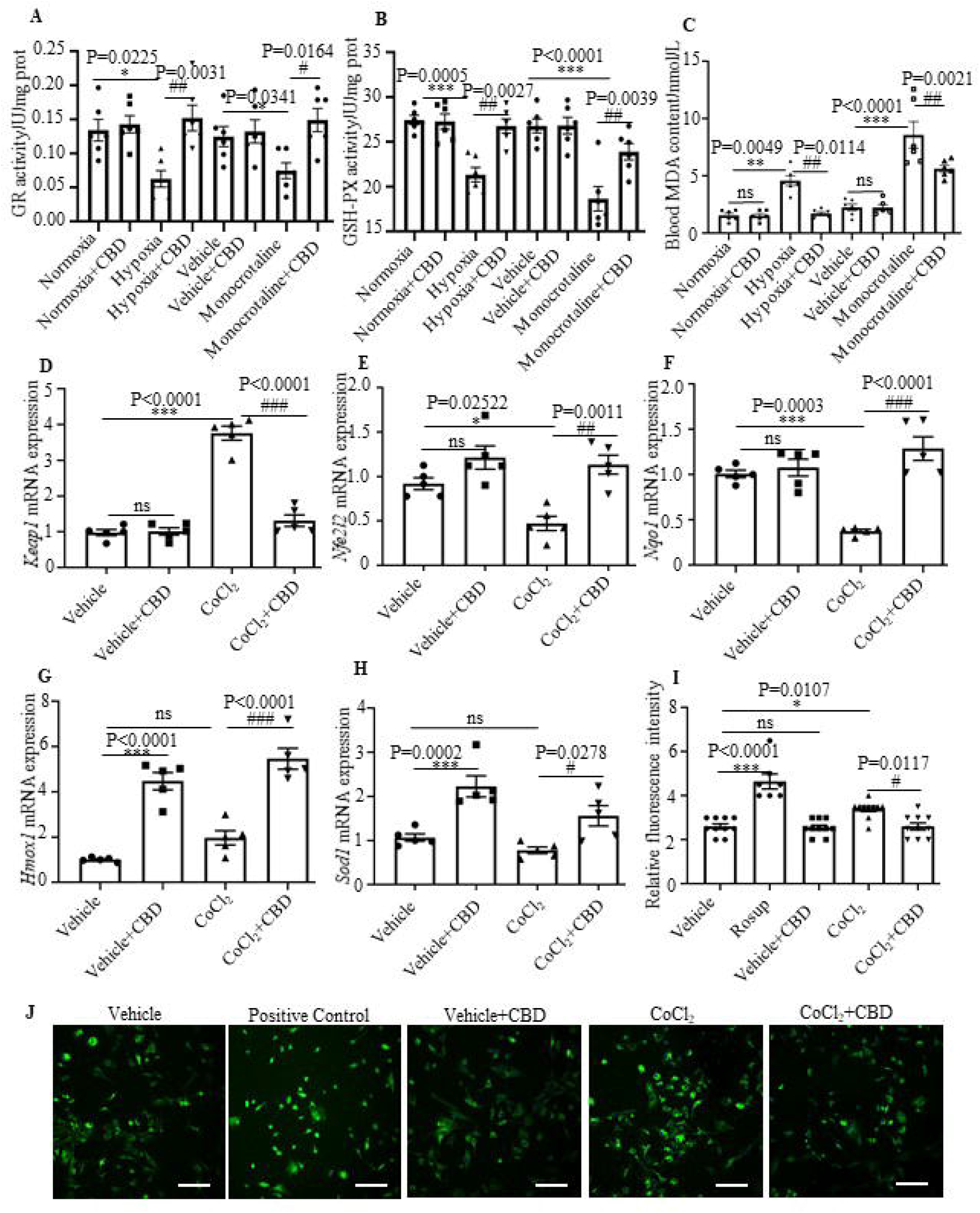
Protection effect of CBD on oxidant stress. **A-B**, Quantification of enzyme activities of GR and GSH-PX in the blood of hypoxia- and MCT-induced PAH mice models, n=6. **C**, MDA content in the blood of PAH mice, n=6. **D-H**, Expression of *Keap1, Nfe2l2, Nqo1, Hmox1* and *Sod1* in mice PASMCs were quantified using RT-PCR, n=5. **I**, Quantification of ROS with a fluorescence 96-plate by a fluorescence microplate reader and DCFH-DA fluorescence intensity in control human PASMCs and hypoxia treatment human PASMCs after the treatment with CBD (10 µM) or control vehicle or rosup (provided in the kit as a positive control for ROS) for 2 h, n=8. **J**, Representative images of ROS fluorescence assessed by laser scanning microscope in control PASMCs and hypoxia treatment human PASMCs. Nuclei were counterstained using DAPI (4’,6-diamidino-2-phenylindole), n≥7. Scale bars=100 µm. The results were analyzed by one-way ANOVA followed by Bonferroni’s multiple comparison test, \**P* < 0.05, \*\**P*< 0.01, \*\*\**P* < 0.001 vs. the control group, and #*P* < 0.05, ##*P*< 0.01, ###*P* < 0.001 vs. the hypoxia or MCT, or CoCl_2_ treatment group.

**Figure 6.**
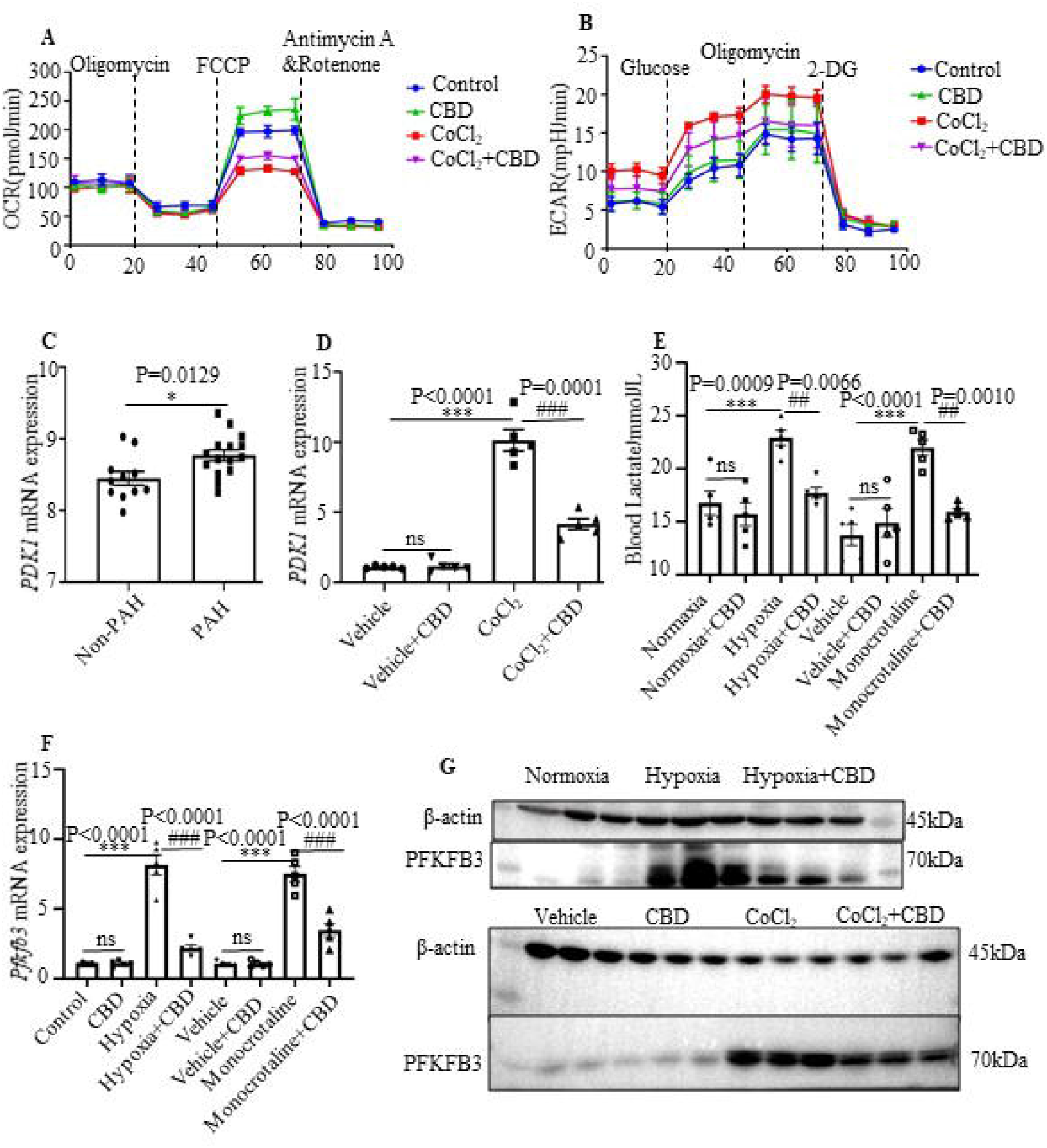
CBD prevented the transition from oxidative phosphorylation to glycolysis in PAH. **A-B**, Quantification of the mitochondrial oxygen consumption rate (OCR) and extracellular acidification rate (ECAR) of control human PASMCs and hypoxia-induced human PASMCs after the treatment with CBD or the control vehicle for 12 h, n=5. **C**, The gene expression of *PDK1* in the lung of patients with PAH (n = 15) and control subjects (n = 11). Values were mean ± S.E.M. and analyzed by an unpaired two-tailed Student’s t-test. **D**, Quantification of *PDK1* in control human PASMCs and CoCl_2_ treatment-human PASMCs after the treatment with CBD or the control vehicle for 12 h, n=5. **E**, Contents of lactate in the blood of hypoxia- and MCT-induced PAH mice were detected with the lactate assay, n=6. **F**, Expression of *Pfkfb3* were quantified in PAH mice lungs with or without CBD treatment, n=5. **G**, Immunoblotting for PFKFB3 in hypoxia-induced PAH mice lung and human PASMCs with or without hypoxia treatment. All blots were re-probed for β-actin as a loading control. The results were analyzed by one-way ANOVA followed by Bonferroni’s multiple comparison test, \**P* < 0.05, \*\**P*< 0.01, \*\*\**P* < 0.001 vs. the control group, and #*P* < 0.05, ##*P*< 0.01, ###*P* < 0.001 vs. the hypoxia or MCT, or CoCl_2_ treatment group.

### 3.5 CBD inhibited glycolysis and reversed the abnormal metabolism in PAH

Oxidative phosphorylation and glycolysis were the main reactions of ATP production, and it had been reported that inhibition of mitochondrial respiration can enhance glycolysis and vice versa [36].We found that CBD treatment reduced the hypoxia-increased glycolysis in human PASMCs (Figure S5E-F). To verify our findings of the glycolysis in vitro and in vivo, we downloaded and re-analyzed the gene expression data of patients with PAH from the public database (No.GSE113439) [37], we obtained the increased expression of pyruvate dehydrogenase lipoamide kinase isozyme 1 (*PDK1*, a key enzyme in glycolysis) in the lung from 15 PAH patients (Figure 6C). Additionally, we observed that CBD significantly reduced the expression level of *PDK1*, which was elevated in hypoxia-induced human PASMCs (Figure 6D). Recently, lactate had been reported to accumulate in PAH patients [38]. And we found that such accumulation of lactate in the blood of hypoxia- and MCT-induced PAH mice models was reduced by CBD treatment (Figure 6E). We then observed that CBD can normalize the high expression level of *Glut1* (Glucose transporter 1) in the lung of PAH mice (Figure S5G). It had been proved that oxidative phosphorylation shifts to glycolysis due to the activation of 6-phosphofructo-2-kinase/fructose-2,6-biphosphatase 3 (PFKFB3) in hyperproliferation of PASMCs in PAH [39]. We observed that CBD could reduce the elevated mRNA level of *PFKFB3* in hypoxia-induced PAH mice models and hypoxia-induced human PASMCs (Figure 6F, Figure S5H), We also obtained the same results of PFKFB3 protein levels in both lungs of PAH mice and in vitro (Figure 6G and Figure S5I). Taken together, our results indicated that CBD can normalize mitochondrial function and inhibit abnormal glycolysis in PAH.

## 4. Discussion

In this study, we demonstrated that CBD elicited the preventive and therapeutic effects in two conventional PAH mice models. Mechanically, CBD could inhibit the hyperproliferation of PASMCs, recovered the dysfunctional mitochondria in hypoxia condition, prevented the lactate accumulation and the excessive glycolysis, and normalized the oxidant stress, which eventually ameliorated PAH, and provided an insight for the novel therapeutic application of CBD in PAH.

In our study, we demonstrated that CBD treatment not only had the drastic efficacy in the preventive model, but it could reverse established disease in two PAH animal models as well. The inhibitory roles of CBD in the elevated RVSP, RVH and the rate of vascular remodeling were remarkable, although the targets or signal pathways of CBD, Bosentan and Beraprost Sodium were different, the comparable efficacy with the first-line PAH drugs also offered possibilities of CBD as a new PAH drug, while as future clinical investigation for the efficacy of the CBD in PAH patients is warrant.

CBD has been reported to elicit its action via several receptors like Cnr1, Cnr2, TRPA1 and PPARγ [40, 41]. In this study, we found that CBD exerted its anti-inflammatory effect in PASMCs independent of Cnr1, TRPA1 and PPARγ signal pathways. And the inhibitors of Cnr2 receptor had the ability to partially counteract the anti-inflammatory effects of CBD in vitro. But no significant phenotypical and pathological differences were observed between the wild type PAH and the Cnr2^-/-^ PAH mice. Moreover, CBD can still reverse the phenotypes of PAH both in WT and Cnr2^-/-^ mice, which strongly suggested that the inhibitory effects of CBD on PAH in vivo is Cnr2-independent. The reasons for this discrepancy are not fully understood yet, we speculated these may due to: 1) in vitro models are routinely used in biological fields for finding signaling targets, while animal models are more relevant to the physiological and pathological conditions; 2) the affinity between Cnr2 and CBD was very low [42], and Cnr2 is not the key mediator for the CBD action in PAH, an unknown novel specific receptor for CBD may exist. In this way, the CBD labeling and CBD-specific receptor fishing experiments are currently under the intensive investigation in our lab.

Generally, the states of fission and fusion maintain a dynamic balance of mitochondria [22]. In our study, we found that CBD could downregulate of fission mediators, notably *DRP1, OPA1*, and upregulate of fusion mediators, especially *MFN1* and *MFN2, MIEF1* both under hypoxic and normoxia conditions in human PASMCs. Accordingly, by using a MitoTracker, we confirmed that CBD treatment increased the proportion of lineage and mean length of mitochondria/pixels of the mitochondrial networks in hypoxia-induced human PASMCs, which resulted in the increased integrity of mitochondrial structure. Whereas how CBD involves in the regulation of each individual gene for the balance of the fusion/fission needs to be further studied.

We also observed that CBD could regulated the genes related to oxidative stress (*Hmox1, Sod1*) in the absence of CoCl_2_, previous studies have also reported the strong antioxidant capacity of CBD, and CBD can upregulate HMOX1 and activate the NRF2-keap1 pathway [34], and normalized the cell functions under the oxidant stress. Based on these results, we speculated that CBD has the efficacy on enhancing mitochondrial functions and maintaining the mitochondrial homeostasis in both path-/physiological conditions.

A recent thorough study from Su’ s group demonstrated that PAH was a condition with abnormal activated PFKFB3 and lactate accumulation in human PASMCs [39]. In line with these studies, we found abnormal activated PFKFB3 and lactate accumulation in PAH mice. In addition, CBD treatment reduced the hypoxia-induced expression of *PFKFB3* and accumulation of lactate in PAH mice and hypoxia-treatment human PASMCs, and subsequently inhibited the transition from aerobic respiration to glycolysis.

CBD is non-psychoactive and with less side-effects [43], which could be further considered as a candidate for new drug investigation, and it may offer a possibility for combined therapy with other known first-line medicines for PAH, but the large scale of clinical investigation on CBD in PAH patients is needed.

## Conclusion

As the Graphic Abstracts showed, we demonstrated that CBD: 1) inhibited the development of PAH via protecting PASMCs from hyperproliferation; 2) restrained the function of mitochondria and alleviate the oxidant stress in PASMCs, which inhibited the excessive glycolysis. Taken together, our study suggested that CBD may provide a promising new insight for the treatment of PAH and other metabolic disorders.

## Abbreviation

PAH: pulmonary artery hypertension
PASMCs: pulmonary artery smooth muscle cells
CBD: cannabidiol
ROS: reactive oxygen species
Nfe2l2: nuclear factor E2 related factor 2
Hmox-1: heme oxygenase 1
Sod: superoxide dismutase
Nqo1: quinone oxidoreductase 1
Cnr: cannabinoids receptor
TRPA1: transient receptor potential A1
PPARγ: peroxisome proliferator-activated receptor
GPR: G Protein-Coupled Receptor
Sugen: vascular endothelial growth factor receptor blocker SU-5416
MCT: monocrotaline
RVSP: right ventricular systolic pressure
RVH: right ventricular hypertrophy
IHC: immunohistochemical
PFKFB3: 6-phosphofructo-2-kinase/fructose-2,6-biphosphatase 3
OCR: O_2_ consumption rate
ECAR: extracellular acidification rate
DRP1: dynamin-1-like protein
FIS1: mitochondrial fission 1 protein
OPA1: optic atrophy type 1 (OPA1)
MFN1/2: mitofusin 1/2
MIEF1: mitochondrial elongation factor 1
GR: glutathione reductase
GSH: glutathione peroxidase
MDA: Malondialdehyde
Keap1: Kelch-like ECH-associated protein 1
Glut1: Glucose transporter 1
PDK1: pyruvate dehydrogenase lipoamide kinase isozyme 1

## Supplementary Materials

Extended Material and Methods section and supplemental figures are available in the Supplementary material online.

## Acknowledgments

We thank Hanma Investment Group Co., Ltd. for providing pharmaceutical and technical support. We also thank professor Zhinan Yin from Jinan University for sharing the Lyz2tm1(cre) Cnr2 knockout mice.

## Funding

This work was supported by the National Key Research and Development Project (2018YFC1004702); Natural Science Foundation of Beijing (20G10613); Natural Science Foundation of China (NSFC31970802); The Project for Extramural Scientists of State Key Laboratory of Agrobiotechnology (2018SKLAB6-19).

## Conflicts of Interest

The authors declare no conflict of interest.

## Notes

### Competing Interest Statement

The authors have declared no competing interest.

