## Supplementary Material for "Cannabidiol Attenuates Pulmonary Arterial Hypertension by Normalizing the Mitochondrial Function in Vascular Smooth Muscle Cells"

**Running title: Cannabidiol and PAH**

[1]State Key Laboratory of Agrobiotechnology, College of Biological Sciences, China Agricultural University, Beijing 100193, China.

[2]Hanma Investment Group Co., Ltd., Beijing, 100010, China.

[3]Yunnan Hempmon Pharmaceuticals Co. Ltd., Beijing, 100010, China.

[4]Department of Nutrition and Health, China Agricultural University, Beijing 100193, China.

[5]Department of Surgery, Chinese PLA General Hospital, Beijing 100071, China.

[*] Xiaohui Lu, Jingyuan Zhang contributed equally to this work.

[#]Correspondence: Xiangdong Li, State Key Laboratory of Agrobiotechnology, College of Biological Sciences, China Agricultural University, Beijing 100193, China,, 86-10-62734389 (Tel/Fax).

**Supplemental Materials**

**Materials and Methods**

The data that support this study are available from the corresponding author upon reasonable request.

**1.1 Chemicals, cell line and kits**

High-purity of CBD, THCV, CBDV, and CFA powders (99.8%) were isolated from economic *Cannabis sativa* (hemp) and provided by Hanyi Biotechnology Beijing Co., Ltd, China, ethanol or DMSO dissolved. Vascular endothelial growth factor receptor blocker SU-5416 (Sugen) and monocrotaline (MCT) were purchased from Sigma, USA. Rimonabant (antagonist of cannabinoids receptor 1, Cnr1, ethanol dissolved), SR144528 (antagonist of Cnr2, DMSO dissolved), AM630 (antagonist of Cnr2, DMSO dissolved), HC030031 (channel blocker of transient receptor potential A1, TRPA1, DMSO dissolved), GW9662 (antagonist of cannabinoids receptor, peroxisome proliferator-activated receptor, PPARγ, DMSO dissolved) were purchased from Sigma, USA. $CoCl_2$ (used for stimulating the hypoxia condition on cells), polysorbate 80, carboxyl methylcellulose sodium and benzyl alcohol were purchased from Sigma, USA. Ketamine and xylazine were purchased from Aladdin, China. Bosentan was purchased from Aladdin, China. Beraprost Sodium was purchased from Sigma, USA.

The kit for MitoTracker mitochondrion-selective probes was purchased from Invitrogen, USA, and ROS assay kit was purchased from Beyotime, China.

The human PASMC cell line (No. BNCC340087) was purchased from Beina biology, China, authorized by ATCC and were cultured in Dulbecco's Modified Eagle's Medium—high glucose (Sigma, USA) according to the supplier's instructions.

**1.2 Animals**

All the experiments were performed in accordance with the NIH guidelines for the Care and Use of Laboratory Animals, and all procedures were approved by the Ethics Committee for Animal Experimentation of the China Agricultural University. C57BL/6J mice purchased from Beijing Vital River Laboratory Animal Technology Co., Ltd. China and housed in a 12 h light/dark cycle under specific pathogen-free conditions. The Lyz2tm1(cre) Cnr2 knockout mice (C57BL/6J background, with Cnr2 knockout in macrophages) were generous gifted from Prof. Zhinan Yin, Jinan University, the Cnr2 knockout mice were generated by hybridized with a tool mice (Dppa3, C57BL/6J$^{\textit{-Dppa3em1(IRES-Cre)Smoc}}$).

**1.3 Sugen-hypoxia induced PAH prevention mice models**

Sugen-hypoxia PAH mice model was induced according to the study previously [1, 2]. Briefly, male C57BL/6J mice (n=10/group) were randomly divided into several groups. Normoxia control (treated with vehicle, 0.1% alcohol, i.g.) and Normoxia CBD (daily treated with 10 mg/kg, i.g.) groups were treated without hypoxia nor Sugen. Hypoxia control (treated with 0.1% alcohol, i.g.) and Hypoxia with CBD (daily treated with 10 mg/kg, i.g.) groups were given weekly subcutaneous injection of Sugen (20 mg/kg) in vehicle (0.4% polysorbate 80, 0.5% carboxyl methylcellulose sodium, 0.9% benzyl alcohol) and placed in an animal incubator CJ-DO2 (oxygen concentration maintained between 9% and 11%, Changsha Changjin Technology Co., China) for 21 days. On the day 22, animals were anaesthetized with ketamine (100 mg/kg) and xylazine (10mg/kg), gradually increasing the concentration within 7 min, the anesthesia depth was monitored with pedal reflex. And right ventricular function was assessed, the hearts, lungs and blood were harvested. The right lung was snap frozen in liquid nitrogen. The left lung was fixed with 4% paraformaldehyde in PBS prior to dehydration and paraffin embedding.

### 1.4 MCT induced PAH prevention mice models

The method for MCT induced PAH was described in the previous study [3]. Male C57BL/6J mice (n=10/group) were randomly divided into four groups. Vehicle group received vehicle (0.1% alcohol, i.g.) alone without MCT, Vehicle and CBD group received CBD (10 mg/kg, i.g.) without MCT. Mice in MCT group receiving MCT (600 mg/kg MCT, subcutaneous injection, weekly), mice in MCT+CBD group received MCT (600 mg/kg MCT, subcutaneous injection, weekly) and CBD (daily treated with 10 mg/kg, i.g.). The inducement lasts for 28 days, all animals were sacrificed on day 29 after the MCT inducement. Tissue harvesting was carried out as described above.

### 1.5 Sugen-hypoxia induced PAH therapeutic mice models

Male C57BL/6J mice (n=10/group) on the therapeutic protocol were given weekly subcutaneous injection of Sugen (20 mg/kg) and placed in an animal incubator CJ-DO2 (oxygen concentration maintained between 9% and 11%), allowed to develop PAH from day 0 to 35. Mice in the nomoxia group and Sugen-hypoxia group were receiving vehicle control (0.1% alcohol, i.g.) from day 22 to sacrifice on day 35. Mice in normoxia-CBD group and Hypoxia-CBD group were treated with CBD (daily 10 mg/kg, i.g.) or vehicle control (0.1% alcohol, i.g.) from day 22 to day 35. All animals were sacrificed on day 36 after the Sugen-hypoxia inducement. Tissue harvesting was carried out as described above for mice.

### 1.6 MCT induced PAH therapeutic mice models

Male C57BL/6J mice (n=10/group) on the therapeutic model were given 600 mg/kg of MCT, allowed to develop PAH from day 0 to 42. Mice in MCT group and MCT+CBD group receiving MCT (600 mg/kg MCT, subcutaneous injection, weekly), while others receiving Vehicle (subcutaneous injection, weekly). Mice in the Vehicle group and MCT group were receiving vehicle control (0.1% alcohol, i.g.) from day 29 to sacrifice on day 42. Mice in Vehicle-CBD group and MCT-CBD group were treated with CBD (daily 10 mg/kg, i.g.) from day 29 to day 42. All animals were sacrificed on day 43 after the MCT inducement. Tissue harvesting was carried out as described above for mice.

### 1.7 Right ventricular hypertrophy and right ventricular systolic pressure

The right ventricular hypertrophy (RVH) was measured by calculating the ratio of the weight of right ventricular (RV) to the weight of left ventricular (LV) plus septum as our study described previously [1].

Right ventricular systolic pressure (RVSP) was measured with a micro-pressure transducer (Samba Preclin 420 LP transducer, Samba Sensors, Sweden). It is an invasive test called heart catheterization, the catheter was insert into the right ventricle of the mouse heart along the right carotid artery, the right ventricular systolic pressure curve and data were measured and analyzed with the BE-EH4 biological signal system (Beijing Baianji Science and technology company, China) as described previously [4].

### 1.8 Immunohistochemistry

For assessment of pulmonary arteriolar muscularization, sections of fixed mouse lung tissue (4 μm) were labeled with monoclonal mouse-anti-mouse/rat/human smooth muscle α-smooth muscle actin (SC-53142, Santa Cruz, USA), followed by mouse enhanced polymer method detection kit (PV-9002, ZSGB-Bio, China). To detect proliferation of PASMCs, the lung tissue sections were also stained for monoclonal mouse-anti-mouse/rat/human Proliferating Cell Nuclear Antigen (PCNA) (SC-56, Santa Cruz, USA) expression. Microscopic images were analyzed using the Olympus BX53 microscope (Olympus, Japan).

**1.9 Vascular remodeling analysis**

After paraffin embedding and sectioning, the lung slides (4 μm thickness) were stained with hematoxylin and eosin (H.E.) for morphological analysis. To assess the degree of pulmonary arterial remodeling, microscopic images of Elastin van Gieson and α-smooth muscle actin–stained lung sections were analyzed using the Olympus BX53 microscope (Olympus, Japan). Pulmonary vascular remodeling was quantified by accessing the medial wall thickness and the percentage of muscularization as descripted previously [5-8]. To determine the degree of medial wall thickness, <50 μm and >50 μm in diameter from each lung were randomly outlined by an observer blinded to mouse group or antibody treatment. The degree of medial wall thickness, expressed as a ratio of medial area to cross sectional area (Media/CSA), were analyzed using image J. A minimum of 20 vessels with diameters ranging from 25 to 75 μm were counted from non-serial lung sections and categorized as either fully, partially or non-muscularized. Statistical significance was assessed by comparing the percentage of fully muscularized vessels between groups. Assessment of muscularization was performed blinded.

**2.0 Isolation of mice PASMCs**

Mice PASMCs were isolated from wild type male C57BL/6J mice and cultured by using a modified method described previously [9]. The purity of PASMCs was checked by IHC analyses with antibody against α-smooth muscle actin (1:1000; Santa Cruz, USA). The purity of mouse PASMCs was ≥90%, which was qualified for further study.

**2.1 Detection of mitochondrial reactive oxygen species**

Human PASMCs were treated with or without 200 μM $CoCl_2$ and/or 10 μM CBD for 2 h, and ROS was detected with Reactive Oxygen Species assay kit (Beyotime, China), the density of fluorescence was normalized with DAPI, which was used to detect all cell nucleus. The fluorescence images were observed using a fluorescence microscope (Nikon, Japan) and quantified with a fluorescence 96-plate by a Fluorescence microplate reader (BioTek, USA).

**2.2 Antioxidative enzyme activities**

The antioxidant enzyme activities of mice whole blood were determined by glutathione reductases (GR) assay kit, glutathione peroxidase (GSH) assay kit and malondialdehyde (MDA) assay kit (Nanjing Jiancheng, China) according to the manufacturer's instructions.

**2.3 Cell death and viability**

Cell viability was measured by CCK8 assay (Beyotime, China).

Cytotoxicity of CBD was assessed by detecting the release of lactate dehydrogenase (LDH) into cell incubation media according to the manufacturer's instructions (Beyotime, China). Normally, the LDH is present in the living cells and will be leak out of the cells once the cells in death. When the LDH content in the cell and in the cell culture fluid were detected spontaneously, we can obtain the relative ratio of live cells and dead cells. The content of the extracellular LDH is used to estimate the cell death rate, the content of the intracellular LDH to estimate the proportion of normal cells, to evaluate the toxicity of the drug concentration on the cells.

Cell proliferation was assessed by BrdU assay. Briefly, cells were incubated with 10 μM BrdU (Aladdin, China) for 20 h and fixed with 4% paraformaldehyde for 30 min at room temperature. After treated with 0.1% Triton-X100 for 15 min, cells were incubated with 2 M HCl for 30 min and

0.1 M sodium borate buffer at pH 8.5 for 10 min. Then, cells were blocked with 5% bovine serum albumin and anti-BrdU antibody (Abclonal, China) at a 1:200 dilution at 4°C overnight. The secondary antibody Goat anti-Mouse IgG (H+L) (ZSGB-BIO, China) was applied for 1h at room temperature. Cell fluorescence images were captured with a fluorescence microscope (Nikon, Japan) and quantified.

**2.4 Mitochondrial morphology detection**

Human PASMCs were treated with/without 200 μM $CoCl_2$ and/or 10 μM CBD for 2 h, then detected with MitoTracker-mitochondrion-selective probes (Invitrogen, USA). Live cell fluorescence images were captured with laser scanning confocal microscope A1 (Nikon, Japan). The nucleus of live cells was stained with Hoechst 33342 (Beyotime, China). Mitochondria from each group were randomly selected and the mitochondrial length was quantified by Image J, the statistical methods were referred from several convincing reports [10-13].

**2.5 Analysis of mitochondrial bioenergetics and cellular glycolysis**

$O_2$ consumption rate (OCR) (mitochondrial stress test) and extracellular acidification rate (ECAR) (glycolysis stress test) were determined by the Seahorse XF 24-3 analyzer (Agilent, USA). Human PASMCs were plated onto cell culture microplates on the day prior to the experiments. Human PASMCs were seeded in 24-well plates (2000 cells/well) in DMEM with 10% FBS treated with vehicle or CBD (10 μM), with or without $CoCl_2$ (200 μM) for 12 h. The cells were incubated at 37°C in a $CO_2$-free XF prep station 60 min before the Seahorse assay to allow the cells to equilibrate with the assay medium. For the mitochondria stress, on the day of the experiment, cells were incubated in XF assay medium (Agilent, USA), supplemented with 25 mM glucose, 1 mM pyruvate for 1 h before the measurement. After the recording of the basal rates of OCR, final concentrations of 1.5 μM oligomycin, 1 μM FCCP and 0.5–0.5 μM rotenone and antimycin A were added (XF Cell Mito Stress Test Kit, Agilent, USA) through the instrument's injection ports. For the glycolysis stress test, after plated and treated human PASMCs properly as described above, 10 mM glucose, 1 μM oligomycin, and 50 mM 2-deoxy-D-glucose (2-DG; glycolysis inhibitor) (XF Cell Glycolysis Stress Test Kit, Agilent, USA) were sequentially injected. Values were normalized by cell staining and counting.

**2.6 Isolation and quantitation of RNA**

Total RNAs were isolated with the TRIzol reagent (Invitrogen, USA). Quantitative real-time RT-PCR (qRT-PCR) was performed on a Light Cycler PCR platform (Roche, USA) in accordance to the manufacturer's instructions. The genes primer pairs were listed in Table S1.

**2.7 Western Blot**

The protein extraction was separated by 12% sodium dodecyl sulfate-polyacrylamide gel electrophoresis and transferred onto the polyvinylidene difluoride membrane (Millipore, USA), the blots were blocked with 2% non-fat milk at room temperature and incubated with antibody for β-actin (Santa Cruz, USA) or PFKFB3 (Abcam, USA), MFN2 (Proteintech, USA) at 4°C overnight. The blots were incubated with secondary antibody Goat Anti-Rabbit IgG (H+L) HRP (ZSGB-BIO, China) for 1h at room temperature. The final exposure was detected with an Enhanced Chemiluminescence Detection kit (Invitrogen, USA) and the density of the bands was analyzed by using Image J (USA) software.

12. Ryan J, Dasgupta A, Huston J, Chen K-H and Archer SL. Mitochondrial dynamics in pulmonary

arterial hypertension. *Journal of molecular medicine (Berlin, Germany)*. 2015;93:229-242.doi:10.1007/s00109-015-1263-5

13. Marsboom G, Toth PT, Ryan JJ, Hong Z, Wu X, Fang YH, Thenappan T, Piao L, Zhang HJ, Pogoriler J, Chen Y, Morrow E, Weir EK, Rehman J and Archer SL. Dynamin-related protein 1-mediated mitochondrial mitotic fission permits hyperproliferation of vascular smooth muscle cells and offers a novel therapeutic target in pulmonary hypertension. *Circulation research*. 2012;110:1484-97.doi:10.1161/circresaha.111.263848

**Supplemental Table**

**Table S1. A summary of the qPCR primer sequence**

| Gene | Source | Forward Primer | Reverse Primer |
|---|---|---|---|
| *Actb* | mouse | GGCTGTATTCCCCTCCATCG | CCAGTTGGTAACAATGCCATGT |
| *Il6* | mouse | TAGTCCTTCCTACCCCAATTTCC | TTGGTCCTTAGCCACTCCTTC |
| *Tnfα* | mouse | CCCTCACACTCAGATCATCTTCT | GCTACGACGTGGGCTACAG |
| *Ccl2* | mouse | TTAAAAACCTGGATCGGAACCAA | GCATTAGCTTCAGATTTACGGGT |
| *Cxcl10* | mouse | CCAAGTGCTGCCGTCATTTTC | GGCTCGCAGGGATGATTTCAA |
| *Hmox1* | mouse | AAGCCGAGAATGCTGAGTTCA | GCCGTGTAGATATGGTACAAGGA |
| *Sod1* | mouse | AACCAGTTGTGTTGTCAGGAC | CCACCATGTTTCTTAGAGTGAGG |
| *Nfe2l2* | mouse | TAGATGACCATGAGTCGCTTGC | GCCAAACTTGCTCCATGTCC |
| *Nqo1* | mouse | AGGATGGGAGGTACTCGAATC | AGGCGTCCTTCCTTATATGCTA |
| *Mfn1* | mouse | CCTACTGCTCCTTCTAACCCA | AGGGACGCCAATCCTGTGA |
| *Mfn2* | mouse | TGACCTGAATTGTGACAAGCTG | AGACTGACTGCCGTATCTGGT |
| *Drp1* | mouse | CAGGAATTGTTACGGTTCCCTAA | CCTGAATTAACTTGTCCCGTGA |
| *Keap1* | mouse | TGCCCCTGTGGTCAAAGTG | GGTTCGGTTACCGTCCTGC |
| *Pfkfb3* | mouse | CCCAGAGCCGGGTACAGAA | GGGGAGTTGGTCAGCTTCG |
| *18s* | human | CTTTGGTCGCTCGCTCCTC | CTGACCGGGTTCCTTTTGAT |
| *PFKFB3* | human | ATTGCGGTTTTCGATGCCAC | GCCACAACTGTAGGGTCGT |
| *HMOX-1* | human | AAGACTGCGTTCCTGCTCAAC | AAAGCCCTACAGCAACTGTCG |
| *SOD1* | human | GGTGGGCCAAAGGATGAAGAG | CCACAAGCCAAACGACTTCC |
| *NFE2L2* | human | TCAGCGACGGAAAGAGTATGA | CCACTGGTTTCTGACTGGATGT |
| *NQO1* | human | GAAGAGCACTGATCGTACTGGC | GGATACTGAAAGTTCGCAGGG |
| *MFN1* | human | GAGGTGCTATCTCGGAGACAC | GCCAATCCCACTAGGGAGAAC |
| *MFN2* | human | CTCTCGATGCAACTCTATCGTC | TCCTGTACGTGTCTTCAAGGAA |
| *DRP1* | human | CTGCCTCAAATCGTCGTAGTG | GAGGTCTCCGGGTGACAATTC |
| *FIS1* | human | GATGACATCCGTAAAGGCATCG | AGAAGACGTAATCCCGCTGTT |
| *OPA1* | human | CGACCCCAATTAAGGACATCC | GCGAGGCTGGTAGCCATATTT |
| *PDK1* | human | CTGTGATACGGATCAGAAACC | TCCACCAAACAATAAAGAGTGC |

| | | | |
|---|---|---|---|
| | | G | T |
| *KEAP1* | human | CTGGAGGATCATACCAAGCAGG | GGATACCCTCAATGGACACCAC |
| *MIEF1* | human | CACGGCCATTGACTTTGTGC | TCGTACATCCGCTTAACTGCC |

**Supplemental Figure**

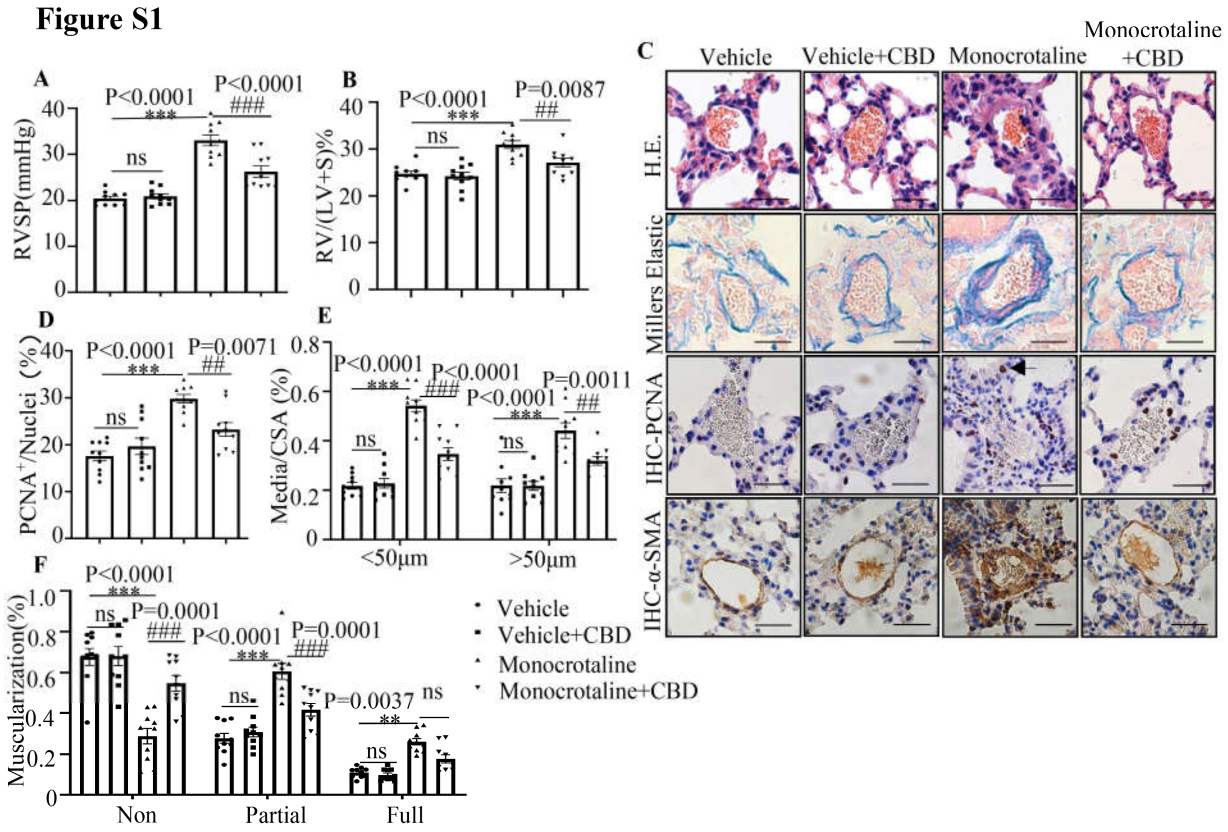


**Figure S1 CBD reversed pathological progression of MCT-induced PAH preventive mice models.**

**A** and **B**, Assessments of RVSP and RVH. **C**, Representative images (×400 magnification) of pulmonary arteries stained with H&E, and representative images (×400 magnification) of vascular remodeling in the distal arterioles stained with elastin or immunostained for PCNA and α-SMA. Pulmonary vascular remodeling rate in the MCT-induced PAH mice, including the quantification of the relative number of PCNA+/nuclei (**D**), the degree of medial wall thickness as a ratio of total vessel size (Media/CSA) (**E**), and the proportion of non-, partially-, or fully muscularized pulmonary arterioles (25 to 75 μm in diameter) from PAH model mice (**F**) (n = 10 mice per group). scale bar=20 μm. Mean +/− SEM. n=10. The results were analyzed by one-way ANOVA followed by Bonferroni's multiple comparison test, *$P$< 0.05, **$P$< 0.01, ***$P$ < 0.001 vs. the control group, and #$P$< 0.05, ##$P$< 0.01, ###$P$ < 0.001 vs. the MCT treatment group.

**Figure S2**

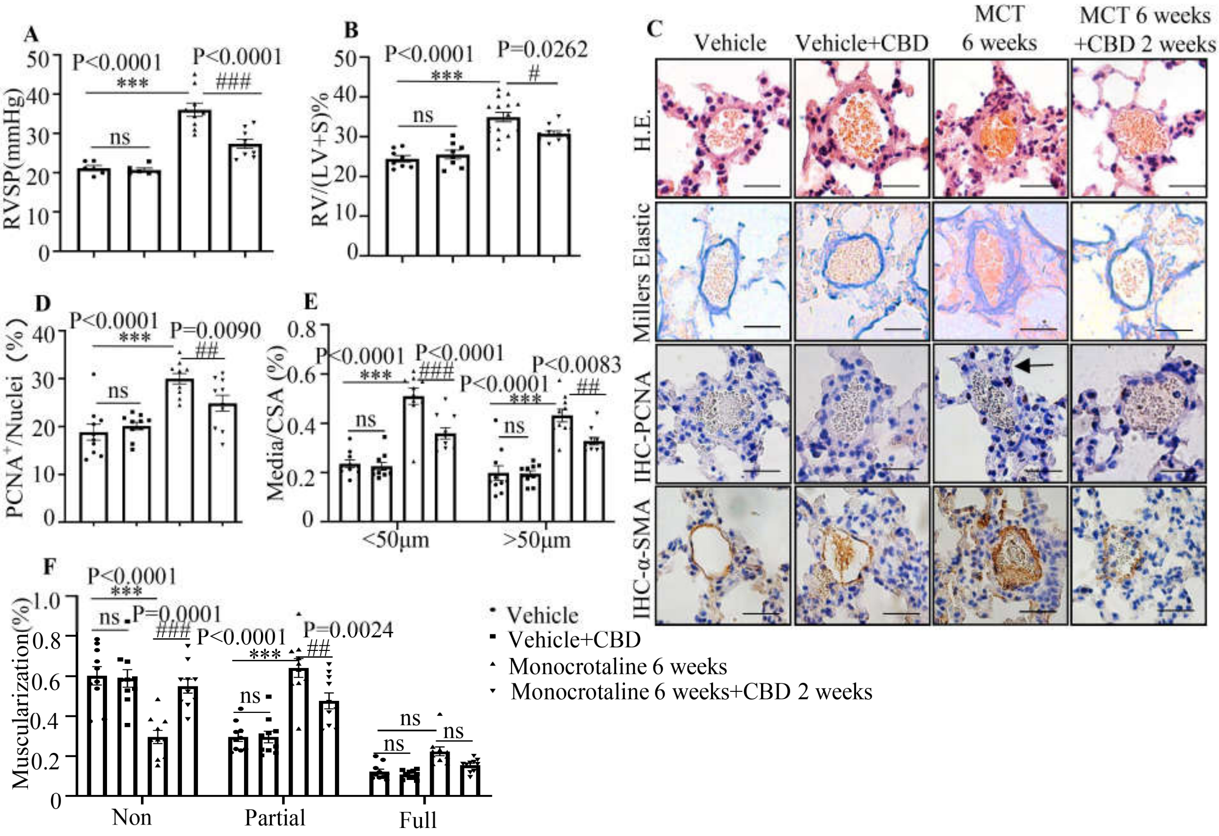


**Figure S2 CBD reversed pathological progression of MCT-induced PAH therapeutic mice models.**

**A** and **B**, Assessments of RVSP and RVH. **C**, Representative images (×400 magnification) of pulmonary arteries stained with H&E, and representative images (×400 magnification) of vascular remodeling in the distal arterioles stained with elastin or immunostained for PCNA and α-SMA. Pulmonary vascular remodeling rate in the MCT PAH mice, including the quantification of the relative number of PCNA+/nuclei (**D**), the degree of medial wall thickness as a ratio of total vessel size (Media/CSA) (**E**), and the proportion of non-, partially-, or fully muscularized pulmonary arterioles (25 to 75 μm in diameter) from PAH model mice (**F**) (n = 10 mice per group). scale bar=20 μm. Mean +/− SEM. n=10. The results were analyzed by one-way ANOVA followed by Bonferroni's multiple comparison test, **P*< 0.05, ***P*< 0.01, ****P* < 0.001 vs. the control group, and #*P*< 0.05, ##*P*< 0.01, ###*P* < 0.001 vs. the MCT treatment group.

**Figure S3**

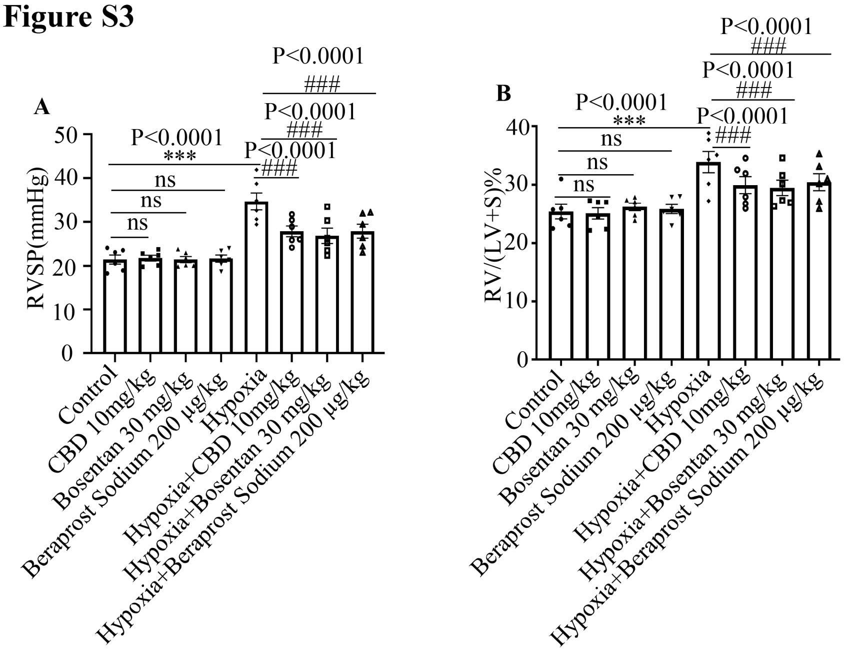


**Figure S3 The compare of the efficacy of CBD, Bosentan and Beraprost Sodium in the hypoxia-induced PAH mice.**

**A and B**, RVSP and RVH of Sugen-hypoxia-induced PAH mice models were assessed, grouped by with or without hypoxia treatment and 10 mg/kg CBD, 30 mg/kg Bosentan daily intragastric administration and 200 µg/kg i.v. Beraprost Sodium once per week for 3 weeks, n=5. The results were analyzed by one-way ANOVA followed by Bonferroni's multiple comparison test, **P*< 0.05, ***P*< 0.01, ****P* < 0.001 vs. the control group, and #*P*< 0.05, ##*P*< 0.01, ###*P* < 0.001 vs. the hypoxia treatment group.

**Figure S4**

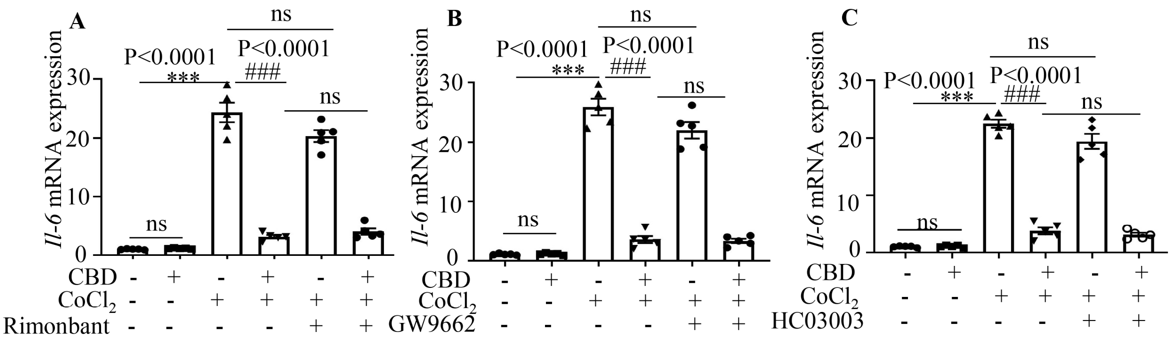


**Figure S4 CBD reduced the expression of *Il6* without the participation of several cannabinoids' receptors in mice PASMCs.**

**A-C**, Expression of *Il6* treated by $CoCl_2$, and the effect of antagonists or channel blockers of CBD receptors (Rimonabant, GW9662 and HC030031) in mice PASMCs, the concentration of them were 10 μM, which were equal to the concertation of CBD, CBD were pre-treated for 30 min with either the antagonists or inhibitors. n=6. The results were analyzed by one-way ANOVA followed by Bonferroni's multiple comparison test, **P*< 0.05, ***P*< 0.01, ****P* < 0.001 vs. the control group, and #*P*< 0.05, ##*P*< 0.01, ###*P* < 0.001 vs. the $CoCl_2$ treatment group.

**Figure S5**

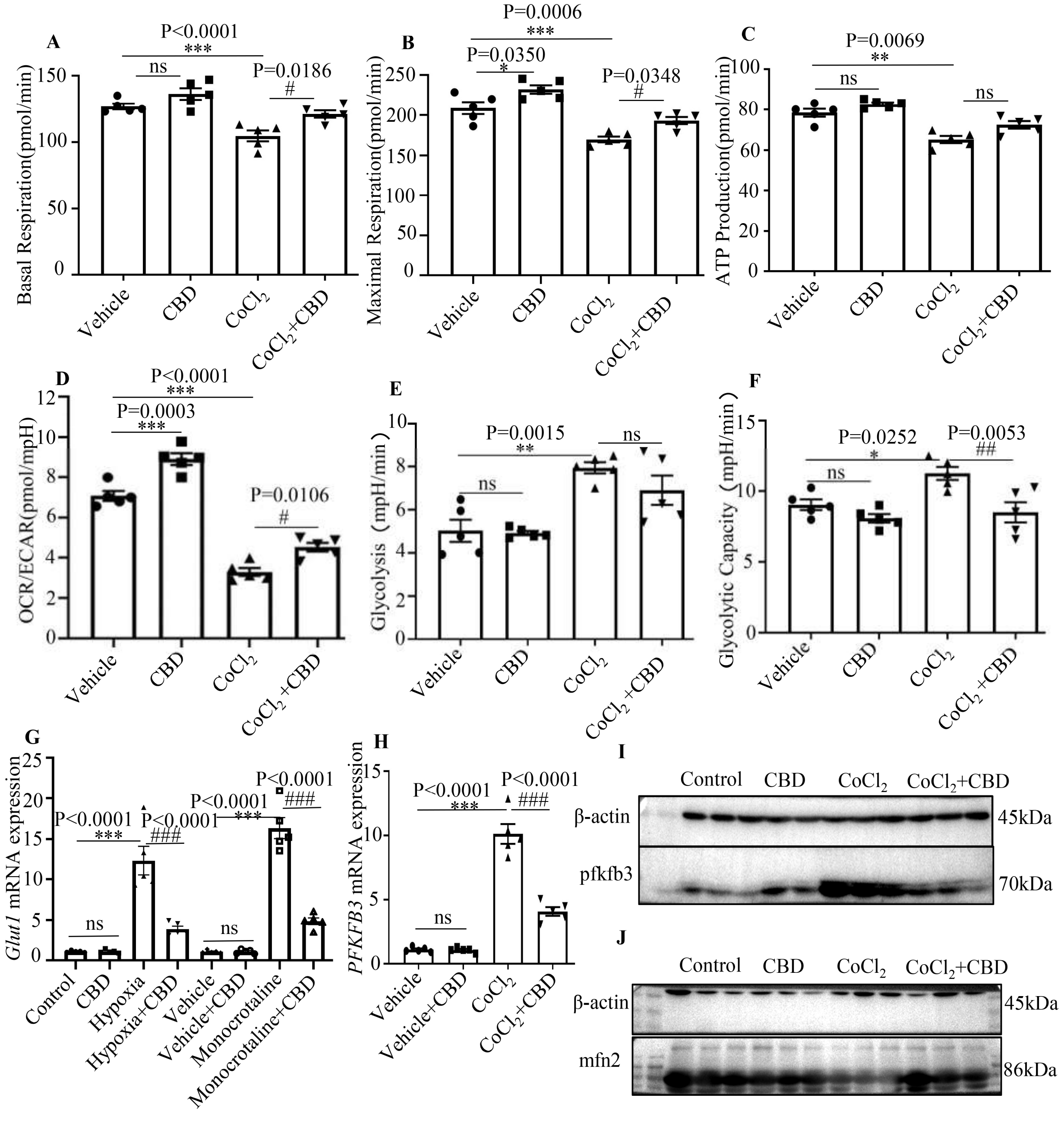


**Figure S5 CBD can reverse hypoxia-induced abnormal glycolysis in both human PASMCs and PAH mice.**

**A-D**, Quantification of the OCR and ECAR in human PASMCs after treatment with CBD or vehicle for 12 hours (n=5 each). Data assessed by mitochondria stress test, including cellular basal respiration, maximal respiration, ATP production and OCR/ECAR. **E** and **F**, Summarized data from glycolytic stress test showing the basal glycolytic rate and the maximal glycolytic capacity. **G**, Expression level of glycolysis marker *Glut1* were quantified in mice lung after inducement for PAH, n=5. **H**, Expression levels of *Pfkfb3* were quantified in human PASMCs with or without $CoCl_2$ and/or CBD treatment, n=5. **I,** Immunoblotting for PFKFB3 in mice PASMCs with or without hypoxia treatment. All blots were re-probed for β-actin as a loading control. **J,** Immunoblotting for MFN2 in human PASMCs with or without hypoxia treatment. All blots were re-probed for β-actin as a loading control. ***$P$<0.001, **$P$<0.01, *$P$<0.05. Mean +/− SEM. The results were analyzed by one-way ANOVA followed by Bonferroni's multiple comparison test, *$P$ < 0.05, **$P$< 0.01, ***$P$ < 0.001 vs. the control group, and #$P$ < 0.05, ##$P$< 0.01, ###$P$ < 0.001 vs. the hypoxia or $CoCl_2$ treatment.
